# Kinetic Model With Feedback Cycle for Age-Dependent Beta Amyloid Accumulation in Mice

**DOI:** 10.1101/2022.10.31.514313

**Authors:** Vivian Tyng, Michael E. Kellman

## Abstract

Amyloid beta (Aβ) is believed to play a key role in Alzheimer’s disease (AD), whose causes, progression, diagnosis, and treatment nonetheless remain poorly understood despite decades of research. Recent studies suggest that Aβ in its various forms participates in multiple mutual feedback loops (“vicious cycles”) including tauopathy, oxidative stress, inflammation, calcium dysregulation, excitotoxicity, and probably many others, eventually leading to neurodegeneration and cognitive decline. Here, we explore a simple kinetic model of a coupled feedback vicious cycle for Aβ buildup based on literature data for Tg2576 mice. The model is used to examine the efficacy of various hypothetical therapeutic approaches, either singly or in combination, to mitigate Aβ buildup. While our computational results support the possible efficacy of combination interventions, they also suggest caution, inasmuch as clear synergy is not found. This kinetic approach highlights the essential importance of a vicious cycle of positive feedbacks in a quantitative model.

## I. INTRODUCTION

There has been a vast amount of research on Alzheimer’s Disease (AD), including its pathogenesis, diagnosis, disease course prediction, and therapeutic interventions. Amyloid beta (Aβ) is a major pathological feature of AD; its key role in the disease [1] is not much disputed. In the amyloid cascade hypothesis [2], Aβ is further proposed as the main cause of AD, even though the related drug development has led to severe disappointment [3]. Current knowledge of the chemical and biological processes involving Aβ remains limited, and a large part of available information has come from *in vitro* studies and animal experiments.

Such data have been collected in animal models, most commonly transgenic mice (although there are growing studies of modified and unmodified higher mammals). In one notable study, Parkinson *et al*. collected the month-by-month data of soluble and insoluble amyloids in the brains of Tg2576 transgenic mice [4]. This and other animal models cannot possibly be fully indicative of the amyloid process in humans [5]. Yet, they offer detailed time-sequence data that are a challenge to explain, and offer a chance to test hypotheses regarding the Aβ process that may help understand similar processes in humans. For the transgenic mice in the aforementioned study, a notable feature is that there is at first a considerable latency period in which the soluble and insoluble Aβ levels are low, followed by a slow but massive (100×) increase terminating at the end of the animal’s lifetime.

In this paper, we explore a nonlinear kinetic model of amyloid accumulation to give an account of observed data in transgenic mice. We first explore the role of nonlinear feedback to give a “vicious cycle,” then consider disease intervention strategies within this kinetic framework. We see this as a step toward investigating similar feedback processes in humans.

An interesting kinetic model for Aβ accumulation that involves highly nonlinear behavior has been proposed by De Caluwé and Dupont [6]. The model hypothesizes a feedback loop between Aβ production and another factor that leads to a vicious cycle. In fact, many such feedback factors have been proposed by various investigators [7, 8], such as tauopathy (excess hyperphosphorylated tau protein, another hallmark of AD), oxidative stress [9], neuroinflammation [10], vascular dysfunction [11], and abnormal glucose metabolism [12]. In this paper we seek a quantitative account of the mice data in Ref. [4] by fitting with a positive feedback kinetic model somewhat similar to that in Ref. [6]. (The two models differ considerably in their presuppositions, dynamical behavior, and motivation, as will be emphasized later at various points.) During the fitting, we find that it is necessary to have a very large feedback to account for the massive increase in Aβ over the mice’s lifetime – whereas pure linear kinetics fails completely.

We get a vicious cycle that gradually but inexorably moves the system from a “healthy” toward a diseased state throughout the mouse’s lifetime. Within the fitting model, we explore possibilities for intervention in the Aβ process by testing hypothetical strategies that affect different kinetic parameters. If they could be implemented in practice, several such strategies could significantly delay or prevent the Aβ level from rising to that observed in mice displaying AD-like symptoms.

It must be emphasized that even within the simple framework of the present models, these mice data are far from providing a definitive understanding of Aβ dynamics. Our goal here is as much to raise necessary questions as to provide even tentative answers. We view the present model as a possible future basis for constructing a model of humans, in which basic aspects of nonlinear kinetics can be used to explore presently unknown aspects of amyloid dynamics. The work here of fitting animal data to a kinetic model is in the spirit of a previous model of autoregulation [13] which subsequently has been used with success to fit experimental data from yeast [14]. (Another approach to modeling autoregulation is found in Ref. [15])

## II. SYSTEM AND KINETIC DATA

Parkinson *et al*. [4] measured the brain concentrations of soluble and insoluble Aβ40 and Aβ42 concentrations in the whole brains of Tg2576 transgenic mice over their lifetime (up to 24 and 26 months for Aβ40 and Aβ42, respectively) by sacrificing the mice at specific ages. In their data, there is a dramatic, lengthy progression of Aβ levels: first a long latency period, then a dramatic rise in the middle (in which the soluble and insoluble fractions increased by a factor of 100 and 1000, respectively), and finally a saturated plateau at later times. Those authors fitted the overall Aβ accumulation by logistic functions. Here, we focus on the soluble Aβ42 because (1) Aβ42 clearly has a much stronger propensity to form the insoluble form than the more common form of Aβ40, and (2) kinetics of soluble Aβ is better characterized than the insoluble form. For the soluble Aβ42, its logistic formula is:

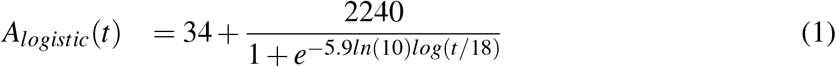

where the Aβ concentration unit is pg/mg, and the time unit is month. In the kinetic models we construct below, the A_*logistic*_ values calculated from Eq. 1 at *t* =1, 2,…, 27 will be used as the target in our kinetic fitting with coupled feedbacks.

## III. KINETIC MODEL

### A. Considerations on kinetics and modeling of mouse data

Before proceeding with a concrete kinetic model, we consider some well-accepted information on the kinetics of Aβ. (1) The turnover time of amyloid monomers (i.e., half-life from synthesis to degradation) is on the order of hours, much shorter than the disease progression months in mice and years in humans) [16]. Therefore, the monomer can be considered in quasi-steady state all the time. (2) According to accumulated animal and human data, the dramatic accumulation of Aβ during disease progression is probably driven by some type of positive feedback process, instead of just linear processes of enhanced production or inhibited clearance [7, 17–19]. For more background about these “vicious cycles” see the Introduction section.

We want to build a model to account for the mice data. First of all, we emphasize that a simple linear model with only steady production of Aβ and its first-order degradation does not reproduce the sigmoidal pattern at all. Specifically, if we take

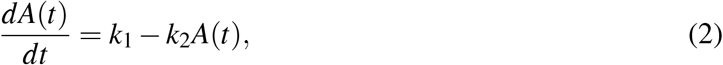

the solution *A*(*t*) = *k*_1_/*k*_2_ + (*A*(1) − *k*_1_/*k*_2_)*e*^−*k*_2_(*t*−1)^ is not sigmoidal (characterized by slow-fast-slow growth rates) for any choices of *k*_1_, *k*_2_, because the growth rate of *A*(*t*) is the fastest at *t* = 0 and decreases continuously in time. So a nonlinear model is needed.

### B. Kinetic model with nonlinear feedback, and fitting to the model

We next suppose that the Aβ participates in “nonlinear feedback cycles” with one or more factors X that interact with Aβ. Considering the simple data, we subsume all feedback factors under one species “X”, whose level varies in time. A diagram of the scheme is shown in Fig. 1.

**FIG. 1:**
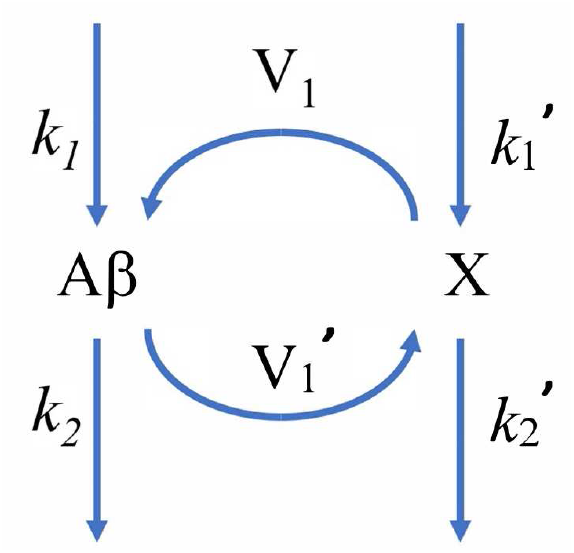
Schematic of the dual-positive feedback model for amyloid (A) and another species X.

In this model, Aβ is produced by two terms, a constant basal rate *k*_1_ and a feedback term dependent on *X*(*t*), and its degradation is first-order with rate constant *k*_2_. X undergoes similar processes including a feedback term driven by Aβ. However, our preliminary fits suggest that the constant basal production rates of Aβ and X, *k*_1_ and 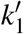 make only inconsequential contributions, and hence they were fixed at zero during the fitting. Thus, we have

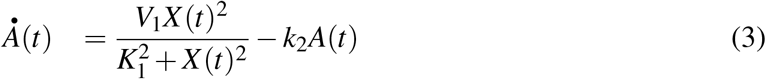

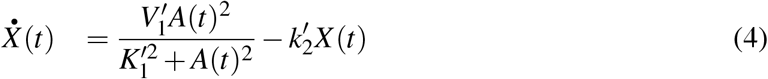

The middle terms in Eqs. 3, 4 are Hill-type functions that reflect the mutual positive feedback between *A* and *X* on each other’s production. *V*_1_ and 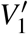 describe the magnitude of the feedback, while *K*_1_ and 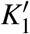 reflect the sensitivity of the feedback. Specifically, 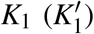 is the *X*(*A*) level when the Hill term reaches half of its maximum, namely 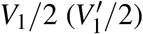 [20]. The Hill coefficient is chosen to be 2, the smallest possible value to produce a sigmoidal trend.

During fitting, parameters in the kinetic model in Eq. 4 were varied to minimize the summed log error in 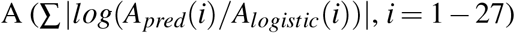, where *A_pred_*(*i*) is the A value predicted by integrating the kinetic equations 3 and 4. *A_logistic_*(*i*) are the values given by the logistic fit in [4] and reproduced here as Eq. 1.

To reduce the number of parameters, we fixed *k*_2_ at 450 (unit: mo^−1^), the value postulated in [4], which comes from the established turnover time of soluble amyloid in mice (1.1 h, compared to 8 h in humans) The initial value *A*(1) = 34 pg/mg is the baseline level obtained experimentally in [4]. For the non-specified X, the initial value was arbitrarily fixed at *X*(1) = 1; our results do not depend significantly on this choice. The fitted parameter values are provided in Table 1, and a comparison of the fit and the target *A_pred_*(*i*) values are plotted in Fig. 2 in both linear and logarithmic scales. It is evident that fitting to the feedback model works well within the limitations of available data.

**TABLE 1:** Kinetic model of soluble Aβ in mouse brain with mutual positive feedback loops. Values indicated with an asterisk are fixed in the fitting.

|  |  |
| --- | --- |
| $k_2$ | 450* |
| $V_1$ | 1.3602E+6 |
| $K_1$ | 10.7556 |
| $A(1)$ | 34* |
| $k_2'$ | 0.00168 |
| $V_1'$ | 0.99795 |
| $K_1'$ | 134.1754 |
| $X(1)$ | 1* |
| LogErr | 0.5397 |

**FIG. 2:**
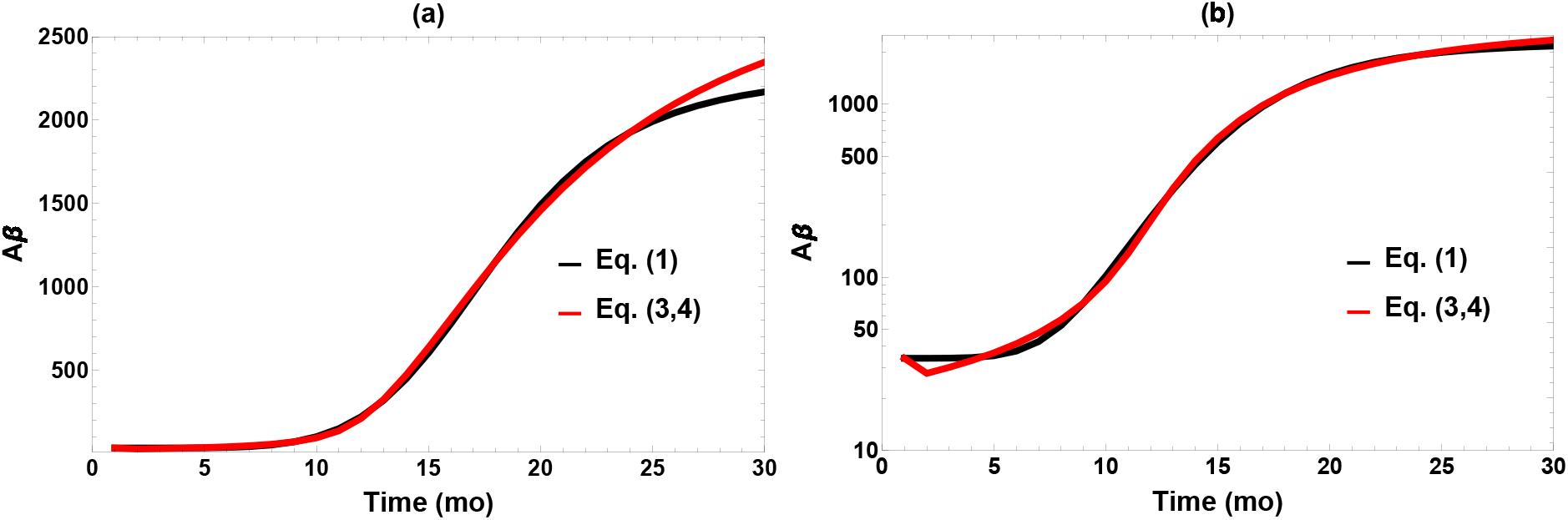
Comparison of fit to the target data (Eq. (1)) plotted in (a) linear and (b) logarithm scales.

### C. The question of bistability

Our vicious cycle model is in some respects similar to the model of De Caluwé and Dupont in Ref. [6], but there are significant differences in the kinetic equations, aims, and presumptions. Their model displays striking features of bistability that play an essential role in their conception of the AD process. They suppose a “jump” from a healthy to a diseased state, associated with a critical point of bistability in their kinetic model. A similar idea of a bistable jump model is outlined schematically by Burlando *et al*. [21]. Does such a picture pertain to the model considered here? This turns out not to be the case, and it is worthwhile to consider why.

The bistable behavior in the model of [6] is suggestive of certain actual features of the AD process in humans. In one study of human subjects not pre-selected for their Aβ status, the brain amyloid levels measured *post mortem* appear to be clustered around a “healthy state” and a “diseased state” [22]. In another longitudinal study of living subjects, the transition from Aβ-negative to Aβ-positive (measured by positron emission tomography) over the years is generally irreversible [23]. These results at least resemble the bistable model of Ref. [6], in which a jump from a healthy to a diseased steady state occurs when a certain bifurcation point of bistability sets in as the Aβ system evolves in time. While highly suggestive, at present the bistable jump model is schematic, qualitative, and speculative, since quantitative time-course data for the brains of living humans are multidimensional, the measurement technologies are far from mature, and the data have intrinsic wide distribution.

We initially were very open to the possibility that our mouse model might show behavior like the jump model of [6]. However, somewhat to our surprise we do not find evidence that bistability plays such a role. The reason is that bistability requires an interaction or competition between linear and feedback processes, whereas the feedback term produced by our fitting to the mice data is massive and overwhelms the linear terms. This moves the “jump-off” bifurcation to an Aβ level way below anything relevant at any point in the mouse life trajectory. Instead, the vicious cycle gradually but inexorably moves the mouse on a lifelong path of disease. This could be called a “drift” picture, rather than a bistable “jump.” It is worth noting that we find this also in variations of our model, e.g., one in which we introduce a linear basal Aβ production term into our kinetic scheme of Eqs. 3, 4. There is simply not a role for the jump, based on the data. It would however be of great interest in the future to have data to tell if the human disease process is more like the present mouse model, or like the jump idea of the model of Ref. [6].

## IV. INTERVENTION IN THE DISEASE PROCESS

Now we use the model developed above to think about strategies for intervening in the mouse disease process, with a view toward eventually using dynamical analysis to devise strategies for AD intervention in humans. Despite decades of extensive search, a safe and effective way to retard AD progression remains elusive. It is now widely thought that the amyloid hypothesis by itself is too simple [24]. In response, there has been a rethinking of concepts of the AD process in recent years. Consistent with this, our model incorporates a feedback loop involving both amyloid and the hypothesized factor “X” to explore possible interventions. Another current line of thought is that single interventions may need to be replaced by combined interventions. We explore this in the context of the model. In the following, we first review tested drug candidates in relation to the principles of the model, then simulate the effect of intervening to vary the different model parameters on the time course of Aβ accumulation. We evaluate the advantages from combinational interventions targeting both linear and nonlinear terms in the model. In the future, we hope to build similar models for humans to explore clinical interventions.

### A. Clinical strategies of intervention

Here we relate strategies of clinical intervention to the somewhat abstract principles of our model. Between 2003 and 2020, no drugs were approved by the US FDA for treating AD although hundreds of substances have undergone clinical evaluations [25]. Aduhelm (aducanumab) broke this trend in 2021, but its approval remains controversial [26]. The development of AD drugs faces many unique challenges, including the disease’s complex etiology, mainly sporadic and late-onset nature, slow progression, a lack of convenient and well-established biomarkers, a shortage of animal models, and the difficulty of drug delivery across the blood-brain barrier. Here, we emphasize three trends relevant to our model. The first is a shift to earlier interventions, preferably before the symptoms appear or even much neurodegeneration has occurred [27]. The second is combining two or more drugs, bioactive substances (e.g., curcumin) [28], or non-pharmaceutical interventions (e.g., physical exercise) together [29]. Examples currently in clinical trial include ALZT-OP1 (ibuprofen and cromolyn, both with anti-inflammation effects) and AMX0035 (tauroursodeoxycholic acid + sodium phenylbutyrate). The third trend is new clinical trial designs that allow researchers to obtain meaningful results sooner and make adjustments during the trial based on new information [30].

The following drug candidates have hypothetical mechanisms relevant to our kinetic model for Aβ:

1. Simply removing Aβ corresponds to increasing *k*_2_ in our model. Quite a few monoclonal antibodies effectively bind to various forms of Aβ and facilitate their degradation: aducanumab (plaques and oligomers), gantenearumab (plaques and oligomers), BAN2401 (protofibrils), solanezumab (monomers), crenezumab (soluble oligomers), and donanemab. Effective removal of Aβ have been observed in phase III clinical trials, yet benefits on the cognitive side remain ambiguous.
2. The production of Aβ depends on various secretases: β-secretase generates Aβ monomer from the amyloid precursor protein (APP), while α- and γ-secreatases catalyze alternative processing of APP to form other products. Several inhibitors of β-secretase have undergone clinical trials, albeit with disappointing results especially in terms of side effects. The drug candidate APH-1105 also enhances α-secretase and thus indirectly suppresses Aβ production. Because our model does not have a basal production of Aβ (by setting *k*_1_ = 0), the effect of β-secretase inhibition would be reflected by reducing the parameter *V*_1_.
3. As mentioned earlier, many biomolecular processes are believed to display a “vicious cycle” with Aβ accumulation [7]. These correspond to species “X” in our developed kinetic model. First of all, many drug candidates target hyperphosphorylated tau, such as TRx0237 (inhibits tau aggregation), ABBV-8E12 (monoclonal antibody), BIIB092 (monoclonal antibody targeting truncated form of tau), and BIIB080 (RNA inhibitor against tau expression). Drugs targeting other vicious cycles include: ALZ-OP1 (anti-inflammation), metformin (targeting insulin insensitivity), and nilvadipine (calcium dysregulation). In our model, the effect of these drugs might be simulated by reducing 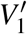, increasing 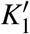, and/or increasing 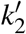.

### B. Effects of single interventions

Now, we discuss possible interventions implemented by changing the parameters in the model. Of the eight parameters in Table 1, *A*(1) and *X*(1) were fixed during fitting, so are not suitable candidates for intervention. Reducing these initial values would not help retard disease progression anyway, considering the massive buildup later. Overall, the accumulation of Aβ could be ameliorated by (1) increasing the parameters *k*_2_ and/or 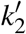, i.e., accelerating first-order removal of *A* and *X*, (2) reducing *V*_1_ and/or 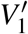, *i.e*., reducing the magnitude of feedbacks, and (3) increasing *K*_1_ and 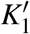 to delay the onset time of feedback without reducing the saturation levels [31]. However, due to a lack of understanding at present of the physiological meaning of *K*_1_ and 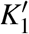, these two parameters are excluded from discussion from this point on. This leaves the four parameters *k*_2_, 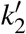, *V*_1_, and 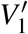 as suitable candidates for intervention. We will first consider single interventions, varying these four parameters separately.

For each intervention, we define the “dose” as

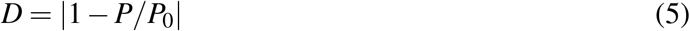

where *P*_0_ and *P* are the kinetic parameter value without intervention (*c.f*. Table 1) and with intervention, respectively. This definition is valid not only for suppressing the production of a biomolecule but also accelerating its removal. In both cases, *D* = 0 means no intervention, and *D* = 1 means doubling the parameter value (for *k*_2_ and 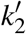) or setting the parameter value to 0 (for *V*_1_ and 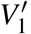). Note that with this definition, a dose *D* > 1 is also possible for the “removal” parameters *k*_2_ and 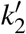, while that of the “production” parameters *V*_1_ and 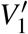 is capped at 1.

Figure 3 displays the outcome of the four single interventions implemented permanently at the age of 1 month (when the mice reach adulthood). Not surprisingly, the Aβ level depends on the dosage for each intervention. At reduced *V*_1_ or 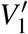, Aβ generally still accumulates albeit at slower rates. For interventions via *V*_1_, 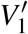, and *k*_2_, a change of −39%, −64%, and +63% (dose = 0.39, 0.64, and 0.63, respectively, according to Eq. 5) would maintain *A*(*t*) ≤ 100 all the way up to *t* = 27. Meanwhile, a comparable change in 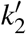 has no noticeable effect on the Aβ buildup (Fig. 3(d)). We see that effective results could be achieved in three of the four single interventions when implemented aggressively, maintaining a high dose for the whole adult life of the mouse. The required minimum dosage is quite sensitive to the specific intervention.

**FIG. 3:**
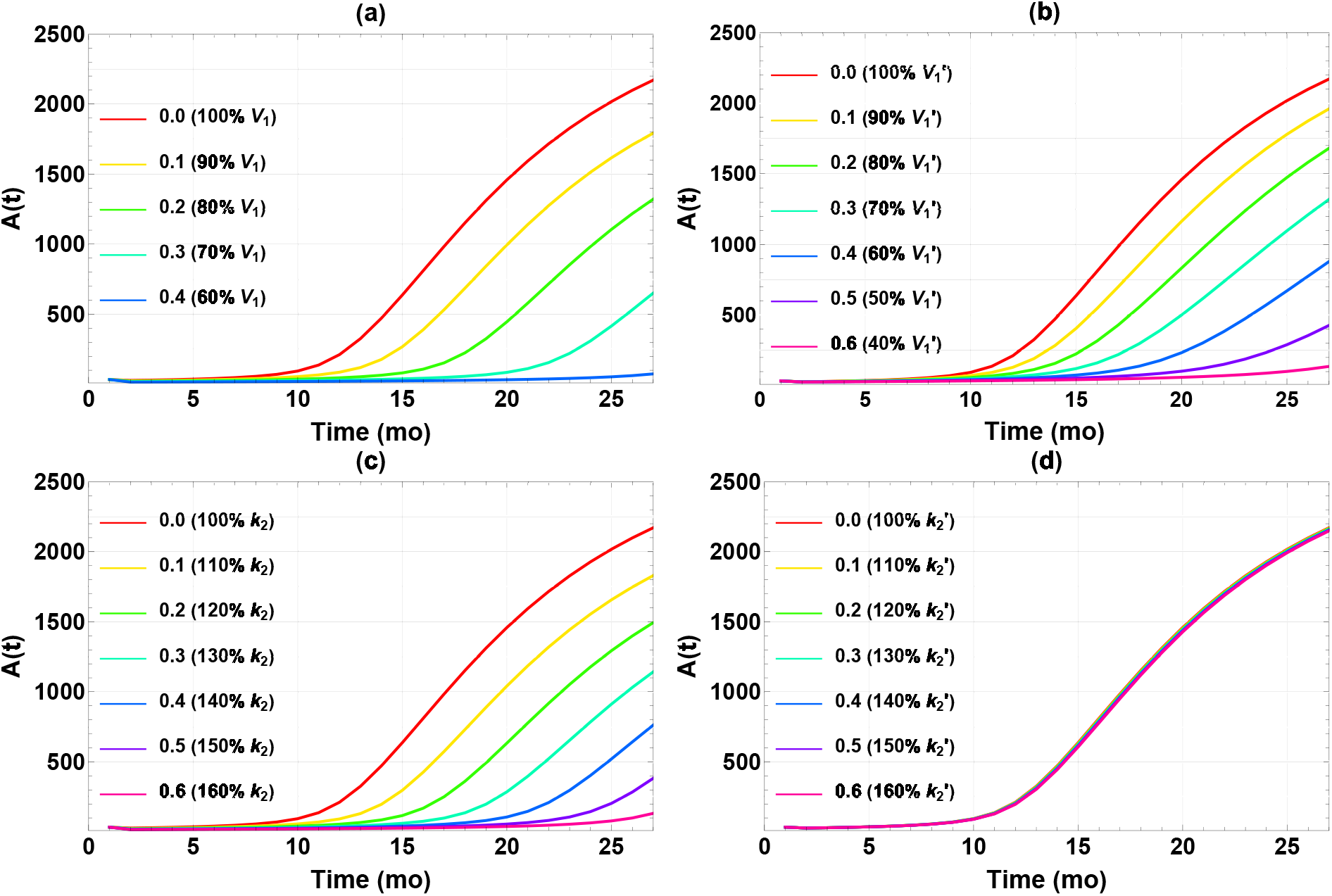
Time courses of *A*(*t*) after implementing different single interventions starting at *t* = 1 mo in (a) V_1_, (b) 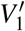, (c) *k*_2_, and (d) 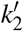. The legends show (left) the doses and (right) the corresponding changes in parameter value relative to those in Table 1.

Figure 4 further illustrates the effect of the single intervention dosage on the final Aβ level (*A*(27)). There seems to be a fairly distinct “threshold” dose for each intervention, above which *A*(27) falls below the target level of 100 and remains low. This is not exactly surprising, since these interventions are designed to reduce *A*(*t*), which is also constrained to be positive. Therefore, *A*(27) falls upon increasing the dosage, even in the less effective case of 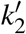 in Fig. 4(d), until the reduction becomes saturated near zero. Still, the thresholds are rather sharp, which has implications for efficacy of interventions. It is essential to get the dose high enough to achieve the target Aβ level of 100. In summary, three of four single interventions work reasonably well if they can be implemented in a high enough dose at the very early age of one month.

**FIG. 4:**
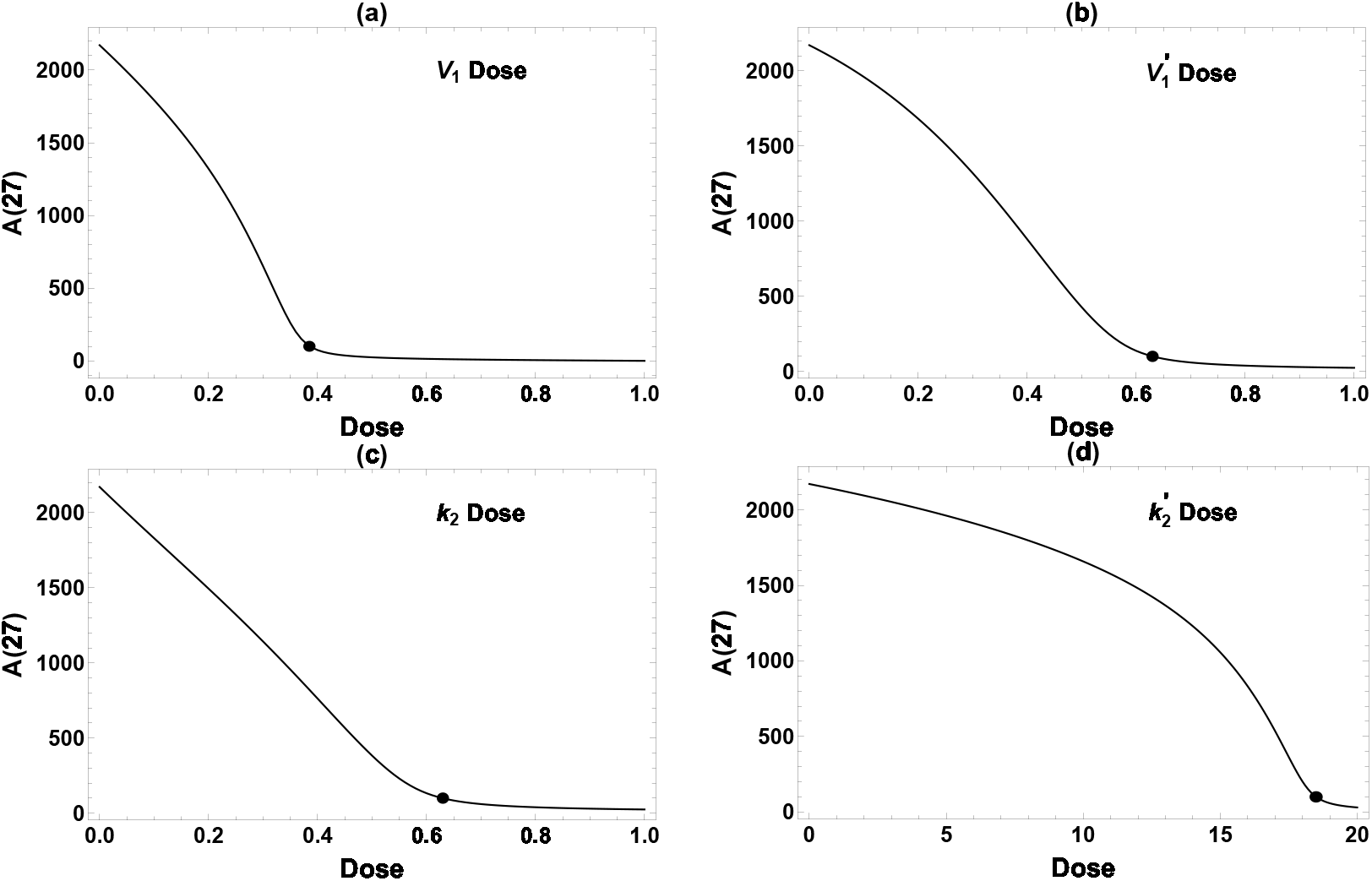
Effect of single intervention dose on *A*(27). All interventions are implemented permanently starting at 1 mo. The dots label the minimum dose required to attain the desired reduction of *A*(27) ≤ 100. Because the model is less sensitive to changes in 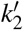, a very high dose is required to achieve meaningful amyloid reduction in (d).

Next, we compared the three single interventions (excluding the ineffective 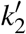) starting at different ages. The corresponding *A*(*t*) data are plotted in Fig. 5. Clearly, the final Aβ level depends on the intervention intensity as well as the timing. A 50% reduction in *V*_1_ (red) is highly effective when starting at *t* = 1, but successful intervention starting at *t* =15 would require a reduction more than 90%. Similar trends are observed for *k*_2_ and 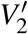. This agrees with the experimental and clinical observations that earlier intervention tends to produce more pronounced effects, although other factors such as side effects and cost-benefit would have to be considered in practice. In the case of Aβ reduction in Tg2576 mice, we conclude that effective intervention would best be implemented at moderate dose and before 5 months, with effective later intervention requiring a much higher, perhaps unrealistic dose. These are rather stringent requirements – starting at 5 months less so than 1 month, but the treatment course still spans most of the lifetime of the mouse. We next turn to the usefulness of combining multiple interventions.

**FIG. 5:**
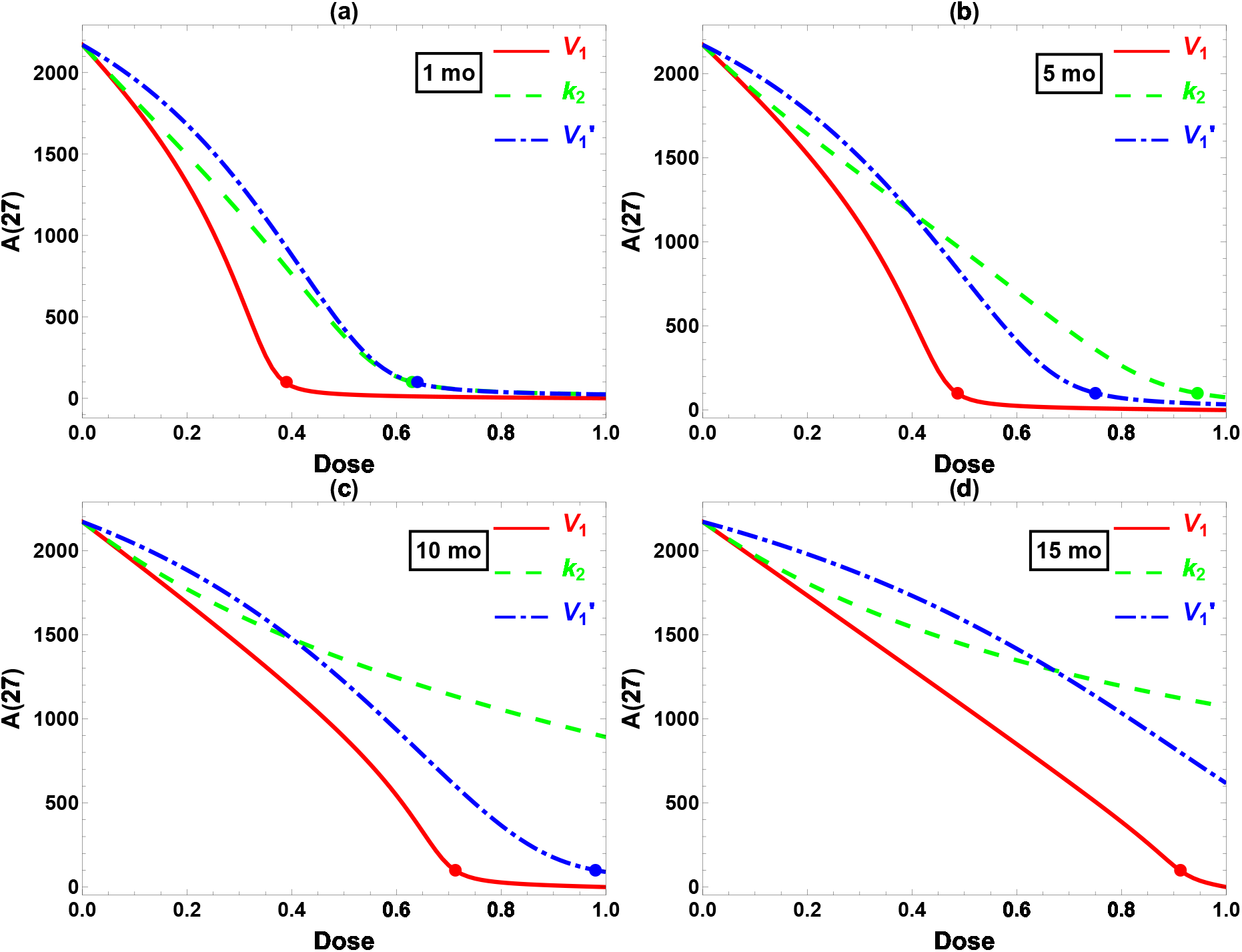
Comparison of interventions in *V*_1_, *k*_2_, and 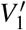 starting at (a) 1, (b) 5, (c) 10, and (d) 15 months. *x*-axis: intervention dose defined in Eq. 5. *y*-axis: Aβ level at 27 months. The dots label the minimum dose required to attain the desired reduction *A*(27) ≤ 100.

### C. Combination Interventions

We have seen that effective single interventions have stringent demands, requiring treatment beginning early in the mouse’s lifespan, and continuing indefinitely. We now consider to what extent multiple interventions offer hope of more effective treatment strategies within the dynamical model. Multi-pronged approaches to treat AD have been advocated for a long time, although only a handful of combined clinical trials have been reported. It is hoped that new trial designs and updated regulatory guidelines [32] would make such trials easier to implement.

Having examined the effect of changing one model parameter at a time, we now combine two parameter changes as interventions. Table 2 shows the dose required to achieve the target *A*(27) < 100 when using either one intervention (diagonal) or a combination of two interventions at the same dose (off-diagonal). (All interventions are implemented permanently after *t* = 1). A smaller minimum dose means more effective treatment. As seen from the table, *V*_1_ monotherapy and all three combinations achieved reasonable efficacy, with a required minimum dose of 0.24–0.39. In contrast, *k*_2_ and 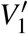 monotherapies have a minimum required dose of 0.63 or higher (as shown earlier in Fig. 3). Clearly, the more effective (single or double) interventions require dampening of the feedback via *V*_1_ and/or 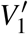, instead of simply accelerating the first-order degradation (*k*_2_, 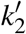). The trick will be to figure out how to intervene on *V*_1_ in the feedback vicious cycle – this seems much less clear than intervening in the linear terms.

**TABLE 2:** Required dose(s) for single and double interventions starting at *t* = 1 to achieve *A*(27) ≤ 100. Diagonal: single intervention. Off-diagonal: two interventions at the same dose (only half of them are shown due to symmetry).

| | $k_2$ | $V_1$ | $V_1'$ |
| --- | --- | --- | --- |
| $k_2$ | 0.63 | 0.24 | 0.34 |
| $V_1$ | - | 0.39 | 0.34 |
| $V_1'$ | - | - | 0.64 |

In Table 2, we combined two interventions at the same strength. However, the doses of individual drugs in actual combinational therapy should be adjusted according to their efficacy and toxicity. Hence, we have further considered two interventions in all possible dose combinations. The results are displayed as 3D plots in Fig. 6. Intersections of the 3D surface with the side walls in the plot correspond to single interventions (*V*_1_, 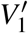, or *k*_2_). The black curve in each panel is the intersection in the horizontal plane of the dose-response surface (orange) and the desired threshold *A*(27) = 100 (light blue). The shortest Cartesian distance *R* between this curve and the origin, indicated by the black line segment from the origin to the black dot, might be taken as the optimal combination. This definition of the optimal dose seems not unreasonable, while certainly being somewhat arbitrary. For present purposes the exact definition should not alter the qualitative conclusions we will present below. (In considering optimality, we ignore the issue of drug toxicity due to the highly abstract nature of our model, although it would be very important when actual drugs are considered.) Among these six plots, the optimal combination generally does not align along the diagonal (*i.e*., combining two interventions at the same dose, as considered in Table 2), highlighting the need to refine dose combinations.

**FIG. 6:**
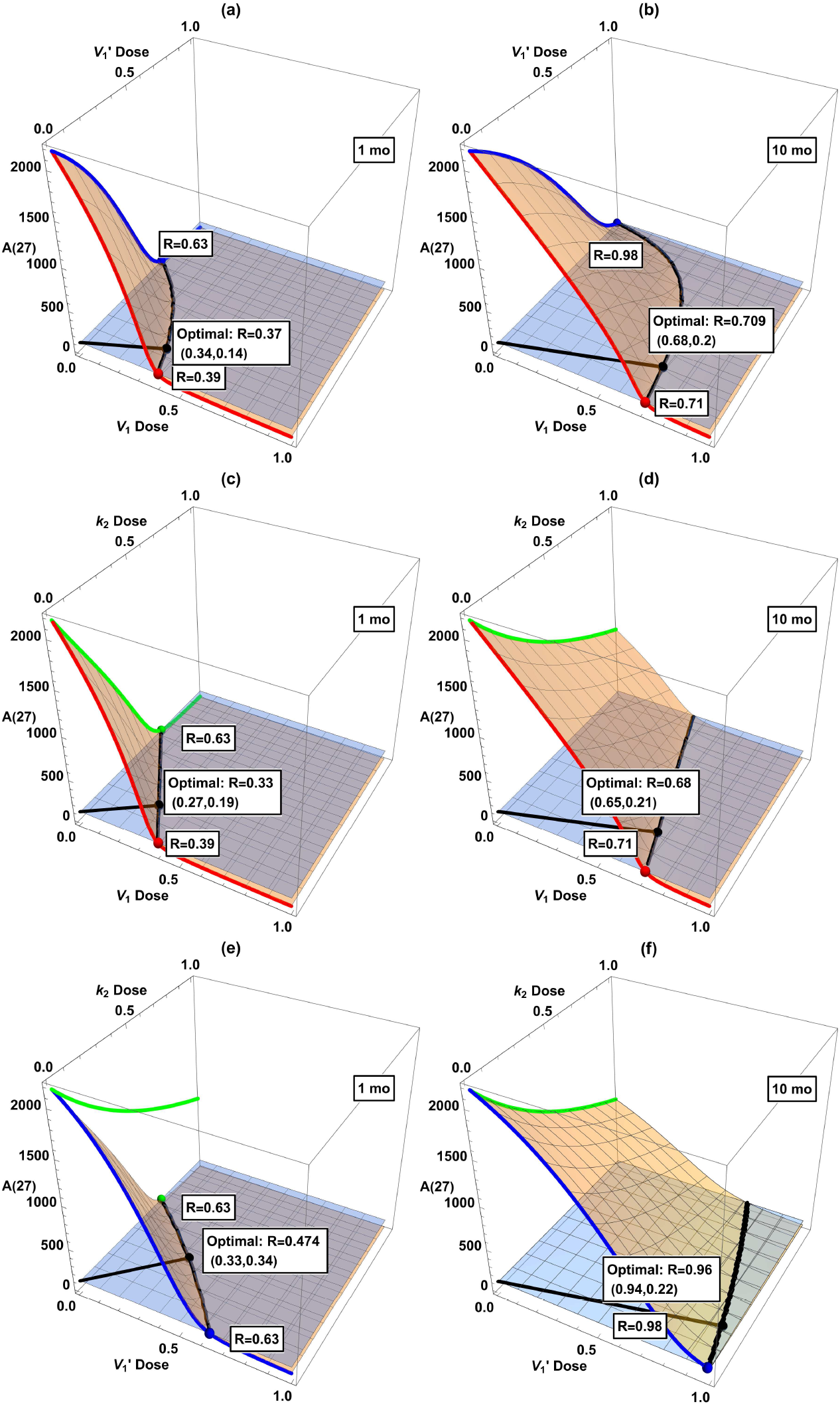
Dose-response surface (orange) when combining two interventions among *V*_1_, 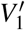, and *k*_2_ starting at 1 or 10 month. The red, green, and blue curves (as well as the dots in matching color) correspond to single treatments in Fig. 5. The horizontal plane (light blue) corresponds to the desired reduction of *A*(27) = 100, and its intersection with the dose-response surface (black curve) indicates the minimum dose combination required to attain *A*(27) ≤ 100. The black dot labels the point with the shortest Cartesian distance from the origin to the black curve of desired reduction (“the optimal combination”).

The dose-response surfaces and optimal dose in dual interventions in Fig. 6 provide information about possible drug synergy. While there is no single quantitative definition of synergy, it broadly means that the combined effect of two separate drugs is larger than that expected from their sum. If the optimal combined doses are substantially smaller than either that of A or B alone, then there is a synergistic effect. On the other hand, if the optimal doses for drugs A+B are each close to that of drug A alone, then the addition of B is hardly useful, let alone synergistic. Unfortunately, the indications from our model suggest that the considered interventions are not especially synergistic in combination. According to Fig. 6, some combinations look favorable if started at 1 mo, e.g. panels (c) and (e). However, the advantage disappears if started later at 10 mo. See panels (d) and (f) with belated treatments, where the optimal dosage is close to one of the axes (*V*_1_ or 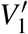 monotherapy), indicating that adding a second intervention in *k*_2_ is hardly useful in these cases. This shows that due consideration to dynamical analysis really is desirable in devising interventions. So far, clinical trials for single interventions of AD have not been terribly promising. Our work here indicates that a cautious attitude should be held toward combinational interventions, at least in the context of the dynamical model of the mice data. Of course, AD pathogenesis is expected to be substantially different between sporadic human cases and genetically engineered mice [33]. To repeat a claim we have made before, it would be desirable to be able to develop a dynamical model based on human data.

## V. CONCLUSIONS

We have proposed a kinetic model with a “vicious cycle” of mutual positive feedback to quantitatively describe reported data of Aβ accumulation in a mouse model. The mouse data and kinetic model display a slow rise in Aβ level at young age, then a lengthy, huge jump (by at least one order of magnitude), and a final slow upward drift towards saturation. The vicious feedback cycle is essential for reproducing the sigmoidal trend in time.

The model has been used to explore various intervention schemes, first single, then double interventions. The results show that interventions must be applied as early as possible. In fact, even strong single interventions fail to achieve meaningful Aβ reduction when implemented beyond a threshold of about 5 months (out of a 27 month lifetime). Similar fairly sharp thresholds exist in intervention strength: an early but insufficiently strong intervention badly fails to suppress Aβ accumulation to a reasonable level. Thus, both the strength and timing of interventions are critical. Between different single interventions, those interfering with the feedback cycle work better than directly accelerating Aβ removal, in terms of the minimum intervention strength required to achieve the same effect. These observations seem consistent with the very limited experimental and clinical observations in humans available so far.

We have also considered various combinations of two interventions. It is encouraging that two interventions of more modest strength can produce comparable results to either intervention at high strength. On the other hand, there is no clear sign of synergy between two interventions. These results may have implications for real interventions [30] in humans.

These key findings reinforce widely held emerging views, especially the importance of early intervention and and also somewhat the advantage of combination treatments. What we have contributed is the crucial role of the feedback cycle, analyzed in a quantitative kinetic model with a dynamical systems approach. Analysis of the model highlights the importance of threshold behavior in the interventions in terms both of the timing and strength. The model brings out the crucial importance of targeting the feedback (which has not been an explicit goal for drug candidates); as well as the efficacy of targeting together both linear removal and nonlinear (feedback) production effects.

In our model, there is a single “X”-factor in the feedback cycle that interacts nonlinearly with the Aβ in the vicious feedback cycle. *tau* protein seems a likely candidate for a possible X. In reality, of course, there may well be many X factors in complex interaction among each other and Aβ. It is important when relevant data become available to further clarify how these various feedback cycles interact. For example, do they act together as if there is effectively one big feedback loop, or is it essentially more complicated?

The positive feedback models proposed earlier by Dupont and De Caluwé [6], and more schematically by Burlando *et al*. [21], similar in some ways to ours, show bistability with a “jump” from a healthy to unhealthy state. This is not the picture seen in our model, which is more like a lifelong steady “drift.” We are focused on a transgenic mouse strain designed to demonstrate extreme Aβ accumulation, and the question of bistability might be different in an examination of analogous human data, if and when they become available. In future work we hope to ascertain data from humans on Aβ with tau as X in the A/X feedback model. If data for both Aβ and tau could be obtained together, it would be possible to apply dynamical analysis fruitfully, with possible insights for intervention.

## Notes

### Competing Interest Statement

The authors have declared no competing interest.

